# Assessing evidence accumulation and rule learning in humans with an online game

**DOI:** 10.1101/2022.02.19.481071

**Authors:** Quan Do, Gary A. Kane, Joseph T. McGuire, Benjamin B. Scott

**Affiliations:** Department of Psychological and Brain Sciences and Center for Systems Neuroscience, Boston University, Boston MA

**Keywords:** decision making, gamification, noise, bias, accumulation, feedback

## Abstract

Evidence accumulation, how the brain integrates sensory information over time, is an essential component of perception and decision making. In humans, evidence accumulation is commonly modeled as a diffusion process in which noise accumulates linearly with the incoming evidence. However, recent studies in rodents have shown that during perceptual decision making, noise scales non-linearly with the strength of accumulated evidence. The question of whether nonlinear noise scaling also holds for humans has been clouded by differences in the methodologies typically used to collect and analyze human and rodent data. For example, whereas humans are typically given explicit instructions in these tasks, rodents are trained using feedback. Therefore, to evaluate how perceptual noise scales with accumulated evidence, we developed an online evidence accumulation game and nonverbal training pipeline for humans inspired by pulse-based evidence accumulation tasks for rodents. Using this game, we collected and analyzed behavioral data from hundreds of participants trained either with an explicit description of the relevant decision rule or merely with experiential feedback. Across all participants, performance was well described by an accumulation process, in which stimuli were integrated equally across time. Participants trained using feedback alone learned the game rules rapidly and used similar strategies to those who received explicit instructions. Decisions in both groups were influenced in similar ways by biases and perceptual noise, suggesting that explicit instructions did not reduce bias or noise in pulse-based accumulation tasks. Finally, by leveraging data across all participants, we show that perceptual noise during evidence accumulation was best described by a non-linear model of noise scaling, consistent with previous animal studies, but inconsistent with diffusion models widely used in human studies. These results challenge the conventional description of humans’ accumulation process and suggest that online games inspired by evidence accumulation tasks provide a valuable large-scale behavioral assessment platform to examine perceptual decision making and learning in humans. In addition, the feedback-based training pipeline developed for this game may be useful for evaluating perceptual decision making in human populations with difficulty following verbal instructions.

**Highlights:**

- Development and validation of an online video game to measure perceptual decision making.
- Humans trained using a feedback-based pipeline exhibit similar strategies and performance compared with those receiving instructions.
- Perceptual noise increases superlinearly with sensory evidence.

## INTRODUCTION

When humans and animals make decisions based on uncertain or incomplete information they can use an evidence accumulation process to integrate independent observations across space and time. Evidence integration is frequently studied in the context of perceptual decision-making in which beliefs or percepts are informed by a temporally extended stream of sensory stimuli. Traditionally, perceptual decision making is modeled by a diffusion process in which noise gradually accumulates together with the evidence. Greater levels of noise tend to increase response times and reduce judgment accuracy (Gold & Shadlen, 2007; Ratcliff & McKoon, 2008). The drift-diffusion model (DDM), used for continuous accumulation tasks, assumes that noise is independent across timepoints, while extensions of this model to pulse-based accumulation tasks have assumed independent noise across individual pulses of evidence (Brunton et al., 2013; Keung et al., 2019). Recent studies in rodents have challenged the assumption of temporally independent noise, demonstrating that perceptual noise, quantified as the variance in the estimated number of pulses of evidence, increases super-linearly with the true number of pulses (Koay et al., 2020; Scott et al., 2015). However, it is unclear whether these results extend to other species as the relationship between perceptual noise and evidence magnitude in humans performing similar tasks has yet to be evaluated.

To date, most studies of perceptual evidence accumulation have utilized psychophysical tasks involving noisy or ambiguous stimuli, such as the random dot kinematograms (RDK) (Julesz, 1971; Stirman et al., 2016) or pulse-based accumulation tasks (Brunton et al., 2013). Human participants in these tasks are typically trained using explicit verbal or written instructions that describe the goals and strategy that should be used to guide decisions. This approach hinders comparison with the behavior of nonhuman animal subjects such as rodents, which learn these tasks through feedback and shaping. Explicit instructions have been shown in several contexts to influence decision strategies and modulate the interpretation of feedback (Ghose & Peterson, 2021; Kirsch, 2021; Palmer et al., 2005). Other work, however, shows evidence for strong parallels between description-based and experiential decision-making tasks (Lukinova et al., 2019).

To provide an alternative platform for further investigations of perceptual decision making in humans, we developed an online video game inspired by visual pulse-based accumulation tasks in rodents (Scott et al., 2015, 2017). Recent studies have used video games as an intriguing alternative to traditional psychophysical tasks (Gesiarz et al., 2019; Spiers et al., 2021; Turkakin et al., 2019). Video games can provide an engaging cognitive training environment for humans and enable data collection on a scale that exceeds what is traditionally feasible in laboratory settings (Chabris, 2017; Gray, 2017; Sibert et al., 2017). Video games can offer a diverse range of timescales, from milliseconds-level responses up to days and even months of extended engagement with exciting narratives. Performance can develop from total novice-hood, in which the players do not know a game’s rules or its victory conditions, to tournament levels of expertises. In addition, video games typically employ feedback-based training, similar to the operant training strategies used in animal studies of perceptual decision making.

In the game we developed, participants controlled an agent in a virtual environment, in which brief, randomly timed flashes delivered from objects in the environment indicated the appropriate path or door for the agent to choose. Following correct trials, feedback was delivered in the form of a change in score and auditory reinforcement. To allow a more direct comparison to animal studies, we developed a training pipeline to allow participants to learn the task without any verbal description of the ideal strategy.

Using this game, we collected behavioral choice data from 971 participants (n=194,200 total trials) who played online. Participants rapidly learned the game through feedback alone and could perform 200 trials in 20 minutes or less. The large number of participants allowed us to fit behavioral models to study how participants’ strategies evolved over time. Analysis of human choice data revealed that humans trained with feedback alone reached peak asymptotic performance rapidly (<=70 trials). The addition of training trials (i.e. trials that were perceptually unambiguous) accelerated the learning rate. In contrast to deterministic feedback, which drove dramatic increases in sensory accumulation, fully random, noncontingent feedback drove a modest decrease in sensory accumulation with task experience and a modest increase in side bias. Participants trained with feedback and training trials showed evidence of having converged on similar strategies to participants who were given explicit instructions.

Behavioral choice was well-fit by a model that incorporated stimulus, bias and reward history information. Behavioral errors were best explained by models that incorporated a form of perceptual noise that scaled nonlinearly with the magnitude of the evidence, consistent with previous studies in rodents (Scott et al., 2015, Koay et al., 2020). At the same time, we observed significant heterogeneity in performance across individuals. This heterogeneity could be captured by using hierarchical Bayesian analysis to fit behavioral parameters to individuals while leveraging data from the large population of participants. Individual fits revealed that even when participants received explicit written instructions on what to do, they exhibited the same perceptual noise scaling, reward history effects, and side bias as those who were trained with feedback alone.

These results challenge core assumptions of the widely used diffusion model of perceptual integration in humans and highlight a potentially useful empirical strategy for evaluating individual differences in evidence accumulation and rule learning. The online, gamified task paradigm is readily scalable to acquire data from large numbers of participants with relative speed and affordability. Furthermore, since participants are trained in a manner that resembles the shaping procedures used in rodent models, this game and training design provide for more direct cross-species comparison. Finally, the game introduces novel opportunities to study perceptual decision making in human participants with difficulty following verbal or written instructions.

## RESULTS

We developed an online game for humans inspired by a pulse-based accumulation task previously designed for rodents (Scott et al., 2015, 2017). Participants used keyboard arrow keys to control an agent in a virtual room (**Figure 1a-d**). The room contained a treasure chest and two doors. The goal of the game was to bring the treasure chest to one of the doors. Briefly flashing stars next to the doors indicated the correct door. If the agent brought the treasure chest to the “correct” door, i.e. the door with the greater number of flashes, a pleasant tone was played along with an increase in a numerically displayed score to signal a reward. All participants played a full session (n=200 trials in a 20 minute period).

**Figure 1:**
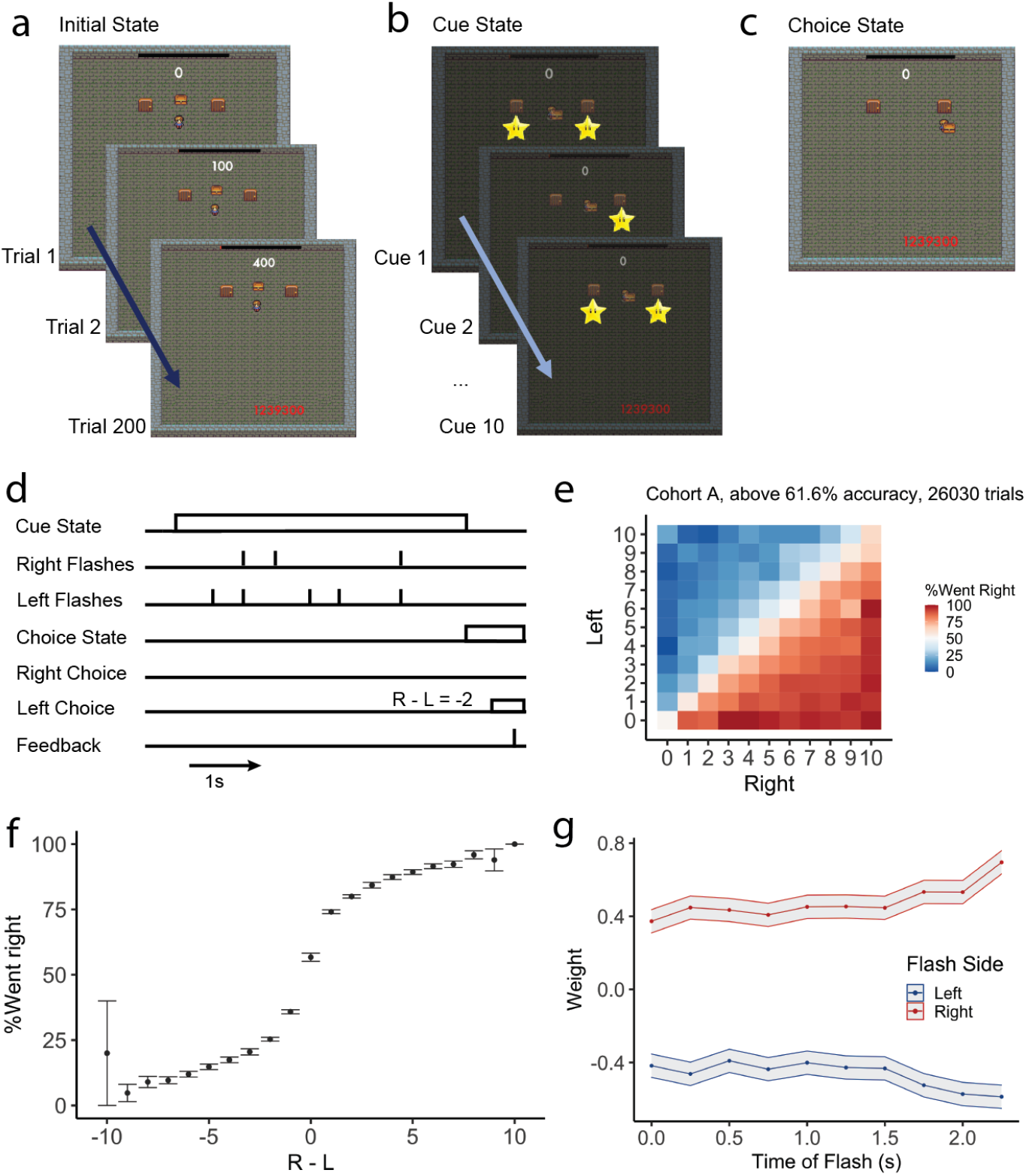
An online game to evaluate evidence accumulation during perceptual decision making. (a) Screen stills from the initial state of trials during the game. At the start of each trial, players were returned to the initial state, and a change in score provided feedback for performance in the previous trial. In this state, the agent was controlled using the keyboard arrow keys. Players entered the cue state after moving the agent and making contact with the treasure chest. (b) Screen stills from the cue state of a single example trial. Flashing stars, located below each door, were displayed as cues. (c) Example still from the choice state, in which players moved the agent to deliver the treasure chest to one of two doors. Delivery of the chest to the door on the side with the greater number of flashes resulted in an increase in the score and auditory feedback. Delivery of the chest to the other door resulted in different auditory feedback. (d) Schematic indicating timing of events during an example trial in which five left flashes and 3 right flashes were presented. (e) Heatmap showing choice data for 137 participants in Cohort A who exceeded 61.6% accuracy, given different numbers of left and right flashes. Color indicates the percent of trials in which participants choose the rightward side. (f) Choice rates as a function of differential evidence strength for the same 137 participants in Cohort A. Dots indicate the mean probabilities of choosing the rightward side. Error bars indicate 95% confidence intervals. (g) Regression analysis showing the relative contribution of left flashes (Blue) and right flashes (Red) occurring at different times in the trial to the participants’ behavioral choice. Lines and error bars indicate the mean and standard error across 137 participants in Cohort A.

### Performance of human participants on the pulse based task

Human performance in this game was first evaluated using a cohort (Cohort A, n=204 participants) that received both written instructions and 10 initial “training trials”. The training trials had large differences in the number of flashes (10 vs 0 and 9 vs 1) to facilitate learning. Performance was evaluated on the subsequent 190 trials.

On average, participants in Cohort A performed the task at 71.4% (+/-14.94% SD) correct. Performance was highly variable across individuals, with some performing near chance (∼50% correct) while others had near perfect performance (>95% correct). We set 61.6% correct as the criterion for inclusion in further analysis – this is the upper bound of the 99% binomial confidence interval for a participant performing at chance. Of the 204 participants in Cohort A, 137 (67%) met this criterion, suggesting that most participants learned to use the flashes to identify the correct side.

Participants who achieved criterion exhibited sensitivity to the relative number of left and right flashes, including differences of a single flash (psychometric slope is significantly different than zero, p<0.001) **(Figure 1e-f)**. Logistic regression analysis of the contribution of individual flashes (See Methods) revealed that participants placed similar weight on early, middle and late flashes to guide their decision **(Figure 1g)**. Together, these results suggest that participants made their decision using an accumulation process in which pulses of sensory evidence were integrated over time to identify the side with the greater number of flashes.

### Effect of explicit instructions and feedback on behavioral strategy

Next we sought to evaluate the effect of explicit instructions on the behavioral strategy used by human participants on this task. We collected data from three additional cohorts (**Table 1**). Participants in Cohort B (n=419) were trained to perform the task using feedback alone, i.e. without a written description of the reward-maximizing decision rule. Like Cohort A, they received 10 initial “training trials”. Participants in Cohort C (n=195) were trained using feedback alone, like Cohort B, but did not experience training trials. Participants in Cohort D (n=153) experienced random feedback, i.e. their score increased on 50% of trials regardless of their response.

**Table 1:** Summary of Cohorts

| Cohort | Number of participants | Training Trials | Task Trials | Instruction | Feedback | Selection Criterion | Number achieving criterion |
| --- | --- | --- | --- | --- | --- | --- | --- |
| A | 204 | 10 | 190 | Yes | Yes | > 61.6% accuracy rate at trial 11 - 200 | 137 |
| B | 419 | 10 | 190 | No | Yes | > 61.6% accuracy rate at trial 11- 200 | 247 |
| C | 195 | 0 | 200 | No | Yes | > 61.6% accuracy rate at trial 11- 200 | 96 |
| D | 153 | 0 | 200 | No | Random | > 61.6% accuracy rate at trial 11- 200 | 0 |

Task performance was comparable between Cohort B and Cohort A (**Figure 2**). A majority of participants in Cohort B surpassed our accuracy criterion (247 out of 417, or 59%, compared to 67% in Cohort A and <1% expected by chance). The difference in the rates at which participants in Cohorts A and B passed criterion was not statistically significant (X-squared = 3.5688, df = 1, p = 0.0589, two-sample test for equality of proportions with continuity correction, alternative hypothesis is two-sided; **Figure 2a**). Among those who reached criterion, we did not observe a significant difference in accuracy (**Figure 2b;** p=0.2356, W = 18156, Mann Whitney U Test) or psychometric function slope (**Figure 2c;** p = 0.1658, W = 18364, Mann Whitney U Test) between Cohorts A and B. Furthermore, flashes in all time bins contributed to choices in both Cohort A and B (**Figure 2d**). Regression analyses indicated that decision weights showed a shallow dependence on time bin that was similar between the two cohorts (linear regression on flash weights on different sides across all time bins; slope = -0.07222, intercept = -0.38367, R-squared = 0.5443, p = 0.0089 for cohort A on the left side; slope = -0.0752, intercept = -0.3384, R-squared = 0.6174, p = 0.0071 for cohort B on the left side; slope = 0.0994, intercept = 0.3655, R-squared = 0.6476, p = 0.003 for cohort A on the right side; slope = 0.0975, intercept = 0.3277, R-squared = 0.6447, p = 0.0032 for cohort B on the right side). Analysis of covariance further suggested there was no significant difference between the two regression lines for Cohorts A and B for flash weights on the left side (Model 1: weight_left ∼ time, Model 2: weight_left ∼ time + cohort, null hypothesis: Model 2 is not significantly better at capturing the data than Model 1, Df= 1, Sum of Sq = 0.0088, F = 4.0971, p=0.06) or flash weights on the right side (Model 1: weight_right ∼ time, model 2: weight_right ∼ time + cohort, null hypothesis: Model 2 is not significantly better at capturing the data than the Model 1, Df = 1, Sum of Sq = 0.008, F = 2.9588, p=0.1036). This suggests that participants trained through feedback alone converged on similar perceptual integration strategies to those participants who received written instructions.

**Figure 2:**
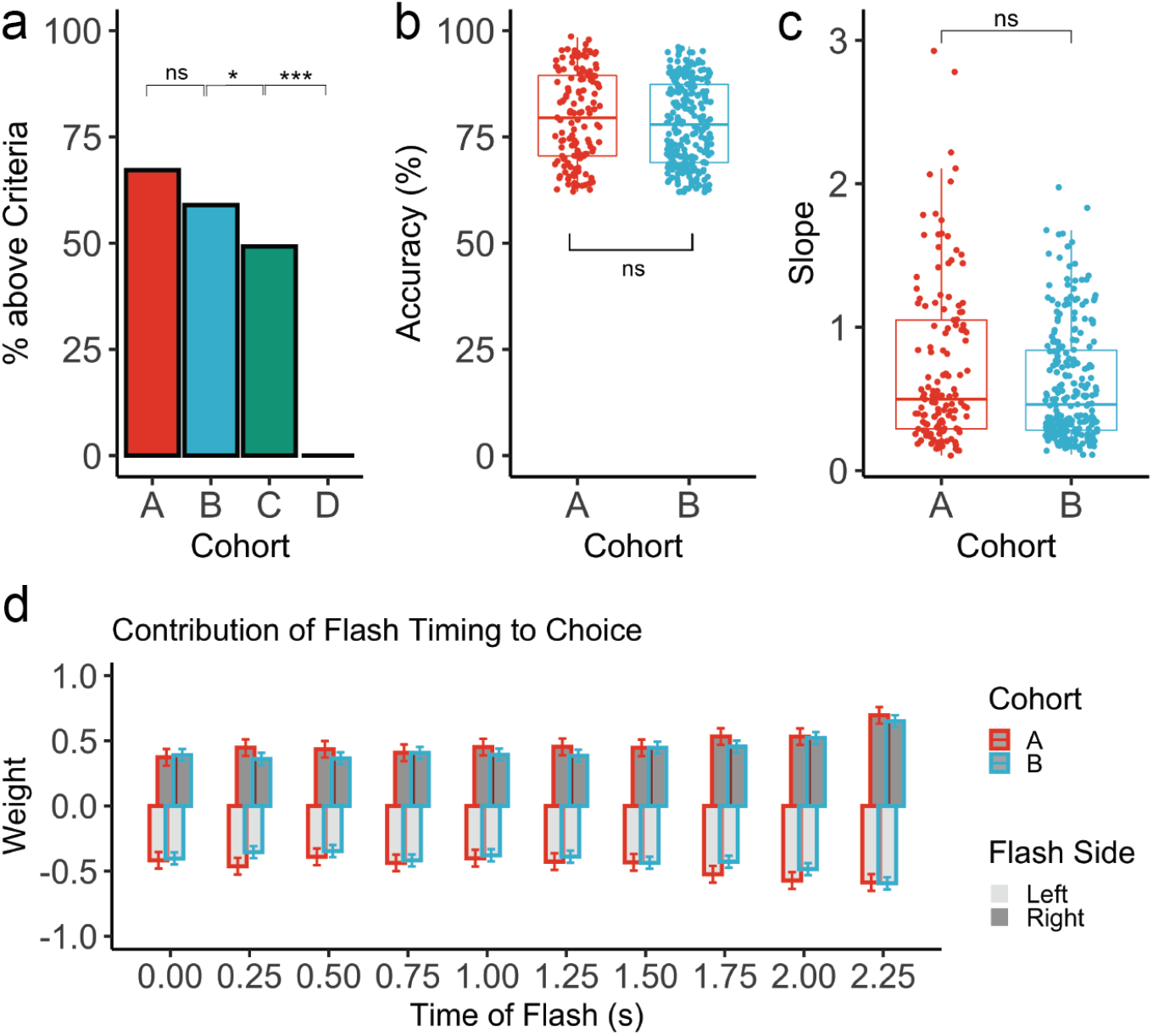
Effect of explicit instructions and feedback on behavioral strategy. (a) Percentage of participants who passed the selection criterion of >61.6% accuracy with 2-sample tests for equality of proportions comparing Cohort A and B (p=0.0589), Cohorts B and C (p=0.03), and Cohorts C and D (p<0.001). (b) Pairwise comparison of accuracy distribution for all participants who passed criterion in Cohort A vs B (p = 0.235) using Mann-Whitney U Test. (c) Pairwise comparison of slope distribution for all participants who passed criterion in Cohort A vs B (p = 0.1658) using Mann-Whitney U Test. (d) Comparing the influence of flashes in each time bin on rightward choices for participants who passed the selection criterion in Cohort A (137 participants) and Cohort B (247 participants). Bar plot indicates the mean estimated weight coefficient for flashes appearing on each side at each time bin. Error bars indicate 95% confidence intervals.

Cohort C (no training trials and no description of the decision rule) exhibited a small but significant reduction in the percentage of individuals who passed the selection criterion (96 of 195, 49%; X-squared = 4.7115, df = 1, p = 0.03, two-sample test for equality of proportions with continuity correction comparing Cohorts B and C), accuracy (W = 14024, p = 0.0085, Mann Whitney U Test comparing all subjects who passed the selection criterion in Cohorts B and C), and psychometric slope (W = 15756, p < 0.001, Mann Whitney U Test comparing all subjects who passed the selection criterion in Cohorts B and C). These results indicate the importance of training trials in allowing participants to rapidly learn the task. Finally, Cohort D (no training trials, no rule description, and random feedback) exhibited a dramatic reduction in the number of participants who reached criterion (0 of 153), accuracy (Mann Whitney U Test comparing all participants in C vs D, W = 26193, p < 0.001), psychometric slope (Mann Whitney U Test comparing all participants in C vs D, W = 19701, p < 0.001). These results further indicate the critical role of feedback in learning the task.

### Effect of feedback on learning and performance over time

Given the critical role of feedback in learning the task, we sought to evaluate how participants altered their performance over the course of the game session. To quantify this change, we aggregated choices across all participants within a cohort and fit the choice behavior in each of 20 10-trial bins to a generalized linear model (GLM) that captured side bias (β_0_), psychometric function slope (β_1_), and reward history effects (β_2_ and β_3_) (**Figure 3a**). We then fit the resulting 20 bin-by-bin coefficients with an exponential function to estimate the learning curve as a function of trial (see Methods).

**Figure 3:**
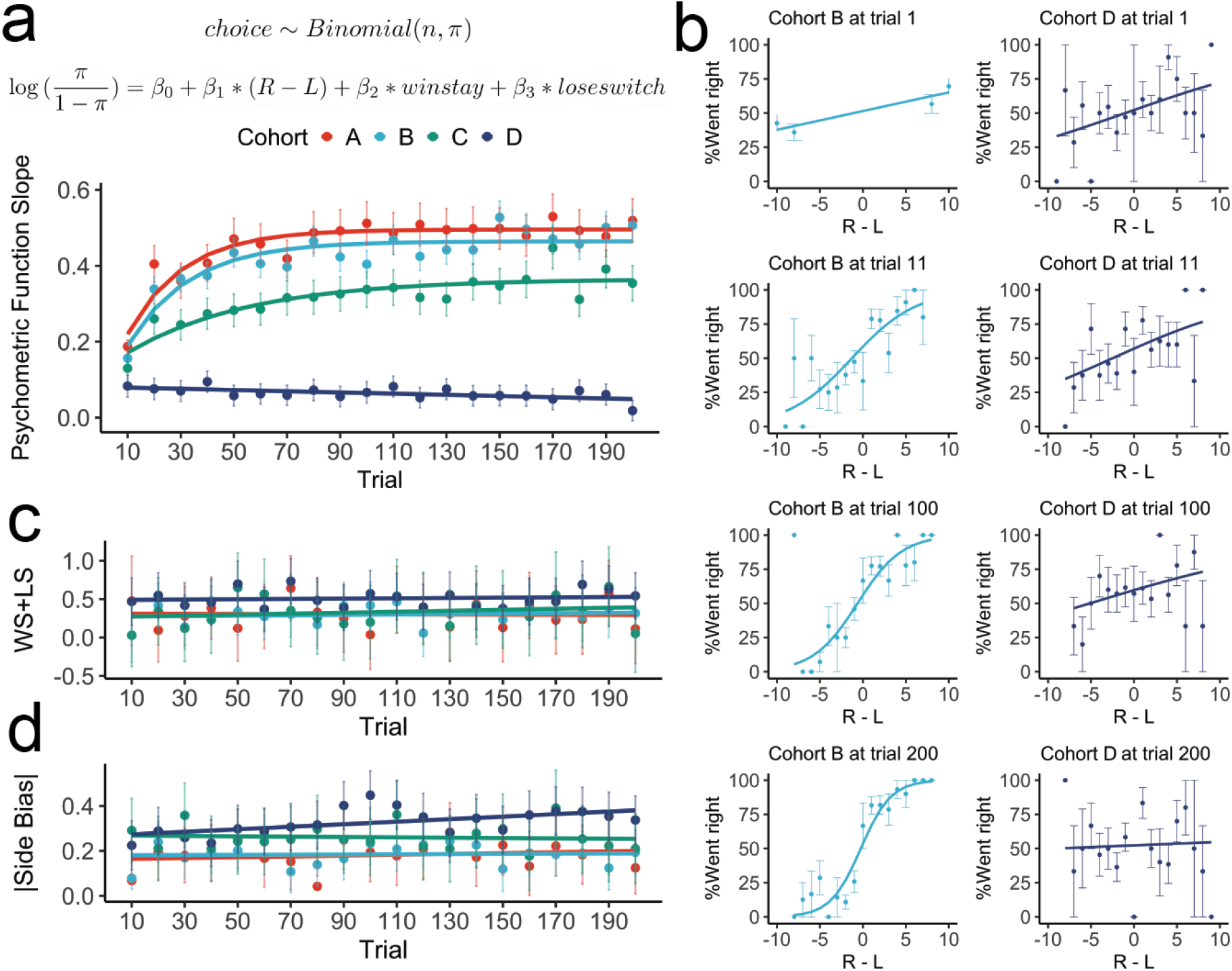
Effect of feedback on learning and performance over time. (a) Choice data from each of the four cohorts was aggregated across each trial and fit to a generalized linear model that incorporated side bias (β_0_), difference in number of flashes (β_1_), and trial history effects (β_2_ and β_3_). Choice was drawn from a binomial distribution for *n* trials with probability π for going to the right side. Analysis was performed on aggregated data from participants who passed the selection criterion in Cohorts A, B and C, and on all participants in Cohort D. Psychometric function slope (β_1_) quantified the contributions of the flashes to the choice within each bin of trials. This parameter was estimated independently in each of twenty successive 10-trial bins. Error bars indicate 95% confidence intervals around the estimated slopes. Learning curves (colored lines) were fit to the psychometric slope (colored dots) in Cohorts A, B and C to quantify changes. A linear model (colored line) was fit to the change in psychometric slope (colored dots) observed in Cohort D. (b) Psychometric plot showing performance change for participants in Cohorts B (247 participants) and D (153 participants) over time, at trial 1, 11, 100 and 200. Participants in cohort B and D both exhibited positive slopes at trial 1. (Note that Cohort B received initial “training trials” so a large right-left difference was always presented on trial 1.) Over the course of successive trials, the slope became steeper for Cohort B but shallower for Cohort D.(c) The win-stay lose-switch parameter was defined to be the sum of β_2_ and β_3_. Error bars indicate 95% confidence intervals around the mean estimated win-stay lose-switch parameter in each 10-trial bin. Linear models (colored lines) were fit to the change in the win-stay lose-switch parameter (colored dots) in Cohorts A, B, C and D. (d) The magnitude (absolute value) of side bias (β_0_) was estimated in each trial bin. Error bars indicate 95% confidence intervals. Linear models (colored lines) were fit to the change in magnitude of side bias parameters (colored dots) in Cohorts A, B, C and D.

All cohorts exhibited significant change in the slope of the psychometric function as they gained experience playing the game. Cohorts A, B and C all had a positive slope (α= 0.473, 0.429 and 0.2149) respectively, suggesting improvement. Interestingly, Cohort D exhibited a small but significant negative slope (α= -0.00159, se=0.00051, p=0.00591), suggesting that their choices gradually became less influenced by the visual cues throughout a game session.

To examine these changes in more detail, instead of binning per 10 trials, we fit the GLM to the choice data for each single trial (bin = 1 trial; 200 bins) using data pooled across participants in Cohort B and, separately, in Cohort D (**Figure 3b**). Consistent with the learning curve analysis, the estimated slope of the psychometric function increased over the course of the session in Cohort B and declined in Cohort D. Both cohorts exhibited a significant positive slope in their psychometric function as early as the first trial (logistic regression, p = 0.0001 for cohort B, p = 0.0271 for cohort D). We point out that these participants indicated that they had not played the game before nor did they receive a verbal description of the decision rule. These results suggest that at least some participants may spontaneously choose the side with the greater number of flashes before receiving feedback.

In addition to the sensory evidence, all cohorts were influenced by reward location history. Participants in Cohort D showed the greatest influence of trial history on choice (linear fit, intercept = 0.5, p<0.001, whereas intercepts < 0.3 for Cohorts A, B and C, p<0.01; **Figure 3c**), suggesting that these participants relied more on reward history to guide choices than other cohorts. The rate of change for the trial history effects was not significantly different from zero for any cohort (all p-values > 0.05), suggesting consistent influence of reward location history over the course of the game session. Finally we computed side bias for each cohort over the game session. On average, Cohorts A, B and C did not show a significant change in the magnitude of side bias over time (regression slope not significantly different from zero). However, Cohort D exhibited a small but significant increase in the magnitude of side bias over the course of the game session (regression slope = 0.00564, se=0.0019, p=0.0089; **Figure 3d**). Together, these results suggest that deterministic feedback drove dramatic increases in sensory accumulation with task experience in Cohorts A, B and C. In contrast, random feedback drove a modest decrease in sensory accumulation and a modest increase in side bias with task experience in Cohort D.

### Perceptual noise scales nonlinearly with the magnitude of the evidence

The drift-diffusion model (DDM), a widely applied model in many perceptual decision making tasks, including pulse-based evidence accumulation tasks in humans, assumes that perceptual noise is independent across timepoints and scales linearly with the amount of evidence. Recent studies in rodents have challenged this assumption, demonstrating that perceptual noise, quantified as the variance in the estimated number of pulses of evidence, increases faster than linearly with the true number of pulses (Scott et al., 2015, Koay et al., 2020). However, it is unclear whether these results extend to other species since the relationship between perceptual noise and evidence magnitude in humans performing similar tasks has yet to be evaluated. Therefore, we used the behavioral choices of participants in our game to evaluate how perceptual noise scaled with evidence (**Figure 4**). We evaluated a family of models inspired by linear and nonlinear noise scaling (**Figure 4a-4b**).

**Figure 4.**
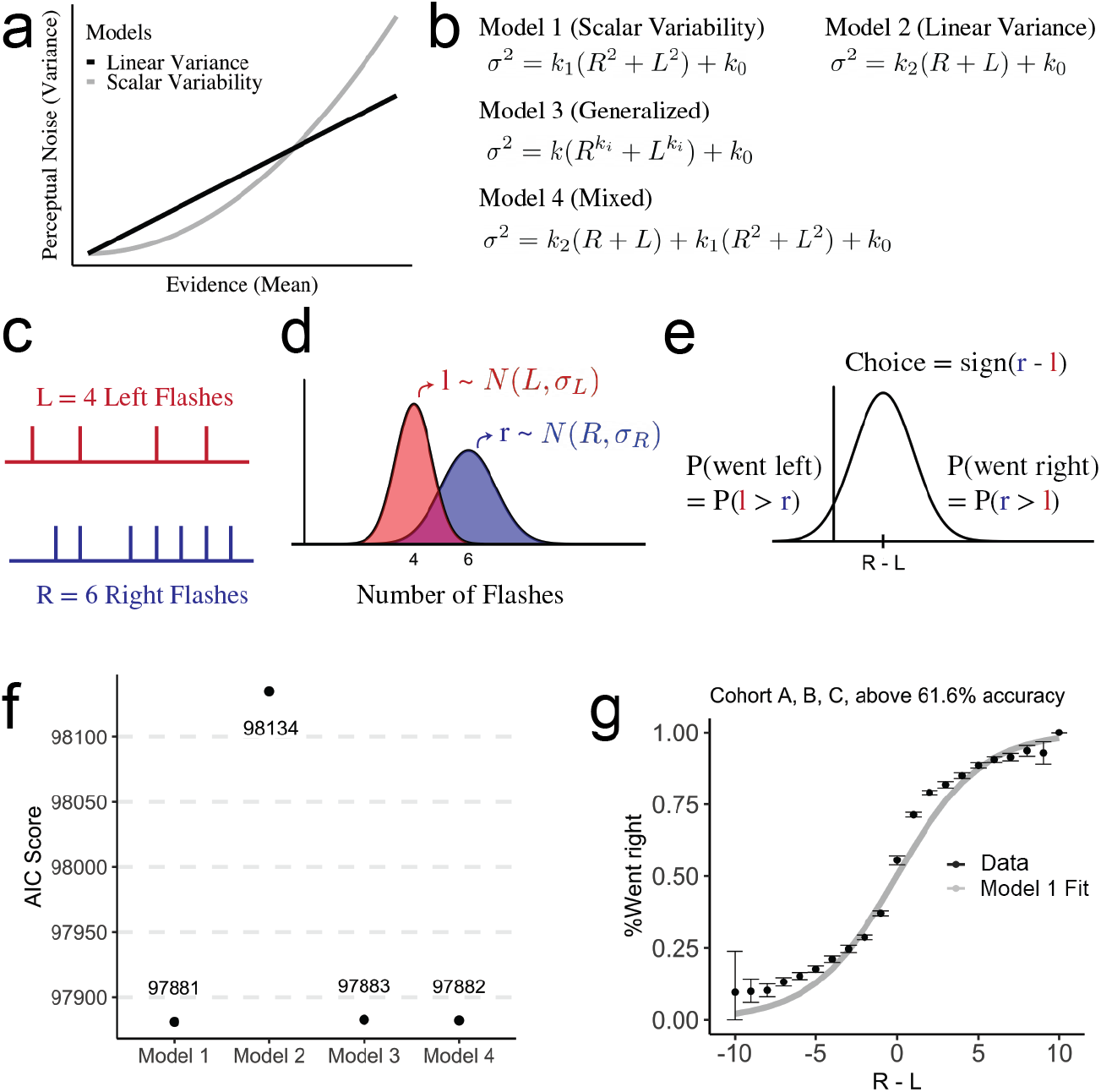
Perceptual noise scales nonlinearly with the magnitude of the evidence. (a) Two different models, linear variance and scalar variability, make different predictions about how perceptual noise scales with the amount of evidence presented. (b) Different models are proposed to test and quantify the prediction made by linear variance and scalar variability. (c-e) Schematic of a signal detection theory-based model to characterize uncertainty in the perception of the number of flashes. (c) Schematic of an example trial indicating the relative timing of the stimuli. In this example trial, 4 flashes were presented on the left side (red) and 6 flashes were presented on the right side (blue). The model assumes that on a given trial, the participant’s estimate of the number of flashes on each side is a continuous random variable drawn from a Gaussian distribution whose mean is the true number of flashes (n) on that side and whose standard deviation is σ. (d) Schematic showing the distributions of perceived number of flashes in the example trial. *l* represents the random variable for the number of left flashes and *r* represents the random variable for the number of right flashes. (e) The choice on each trial is made by sampling and comparing the two random variables *l* and *r*. Error occurs when the difference between the two random variables (greater magnitude -lesser magnitude) is less than zero. (f) AIC scores from a Maximum Likelihood Estimation procedure revealed that the best fitting model was Model 1, which predicts that the standard deviation σ scales linearly with the number of flashes with slope k1 and intercept k0. (f) Psychometric function showing choice data for participants who exceeded a 61.6% accuracy rate in Cohorts A, B and C (black dots and error bars) plotted alongside the prediction made by model 1 (gray line).

We designed four different models that differed in the relationship between perceptual noise (variance of a Gaussian) and number of flashes (mean of the Gaussian). In Model 1, perceptual noise scaled with the number of flashes squared. In Model 2, perceptual noise scaled linearly with the number of flashes. Model 3 was a generalized model with a free parameter for the exponent relating perceptual noise to the number of flashes. Model 4 was an additive mixture model that incorporated both Model 1 and Model 2, with free parameters for the mixing weight of each model (**Figure 4b**).

These models were based on a signal detection theory framework that assumes the participant’s estimate of the number of flashes presented to each side is a random variable drawn from a Gaussian distribution. The mean of the Gaussian is the true number of flashes on that side and the variance or perceptual noise is controlled by free parameters in the model (**Figure 4c-e**). The difference of those Gaussians predicts the participants’ performance: correct trials occur when the difference of Gaussians is positive (i.e. the random variable drawn from the distribution representing the larger number is in fact larger than the random variable drawn from the distribution representing the smaller number) (**Figure 4e**).

We then used maximum likelihood estimation to fit each model to the aggregated choice data from participants in Cohorts A, B and C who passed the selection criterion. The returned AIC scores indicated that Model 1 was the best-fitting model (**Figure 4f**). Models 3 and 4 returned a similar log-likelihood but were penalized for the number of parameters (Models 1 and 2 had 2 free parameters while Models 3 and 4 had 3 free parameters). The best-fitting parameter values for Models 3 and 4 provided further support for the predictions made by Model 1 (exponent k_i_ of 1.951 returned by Model 3, and greater weight placed on the scalar variability term, 1.5 for k_1_ versus 0.289 for k_2_, returned by Model 4). Model 1 provided a good fit to the behavioral choice data from Cohorts A, B, and C (n = 487) (**Figure 4g**). On the other hand, not only did Model 2 return the worst fit based on the AIC score (**Figure 4f**), Model 2 also placed negligible weight on the k_2_ parameter (k_2_ = 1.71e-06), suggesting minimal contribution of diffusion noise to the decision process. We repeated the analysis excluding the first 30 trials across participants in cohort A, B and C to reduce the influence of learning dynamics (**Figure 3a**). This analysis yielded similar findings, with scalar variability still the best-fitting model (**Supplementary Figure 2**). Together, these results suggest that noise scaled nonlinearly with the number of pulses of evidence in humans, similar to the pattern seen in rodents.

### Hierarchical Bayesian Model reveals heterogeneity of noise and bias parameters across individuals

A useful feature of the game-based task described here is the ability to collect data from large numbers of participants online (n>900 in the current study). Such an approach could be used to evaluate participant-to-participant variability in decision making across the population. To compare decision-making strategies across individuals who learned the task (n=487 above criterion), we implemented a Hierarchical Bayesian Model (HBM) to retrieve parameters from the signal detection theory framework for each of the participants. In this model we fit the signal detection theory model, described above, at both the population level and the individual level (**Figure 5a, Supplementary Figure 3**). This modeling approach allowed us to leverage data from a large number of participants to achieve good fits to individual data even with a limited number of trials per individual (200 trials in the current study). Fitting this model to simulated human choice data revealed that this approach was sufficient to accurately recover generative parameters (**Supplementary Figure 4**).

**Figure 5.**
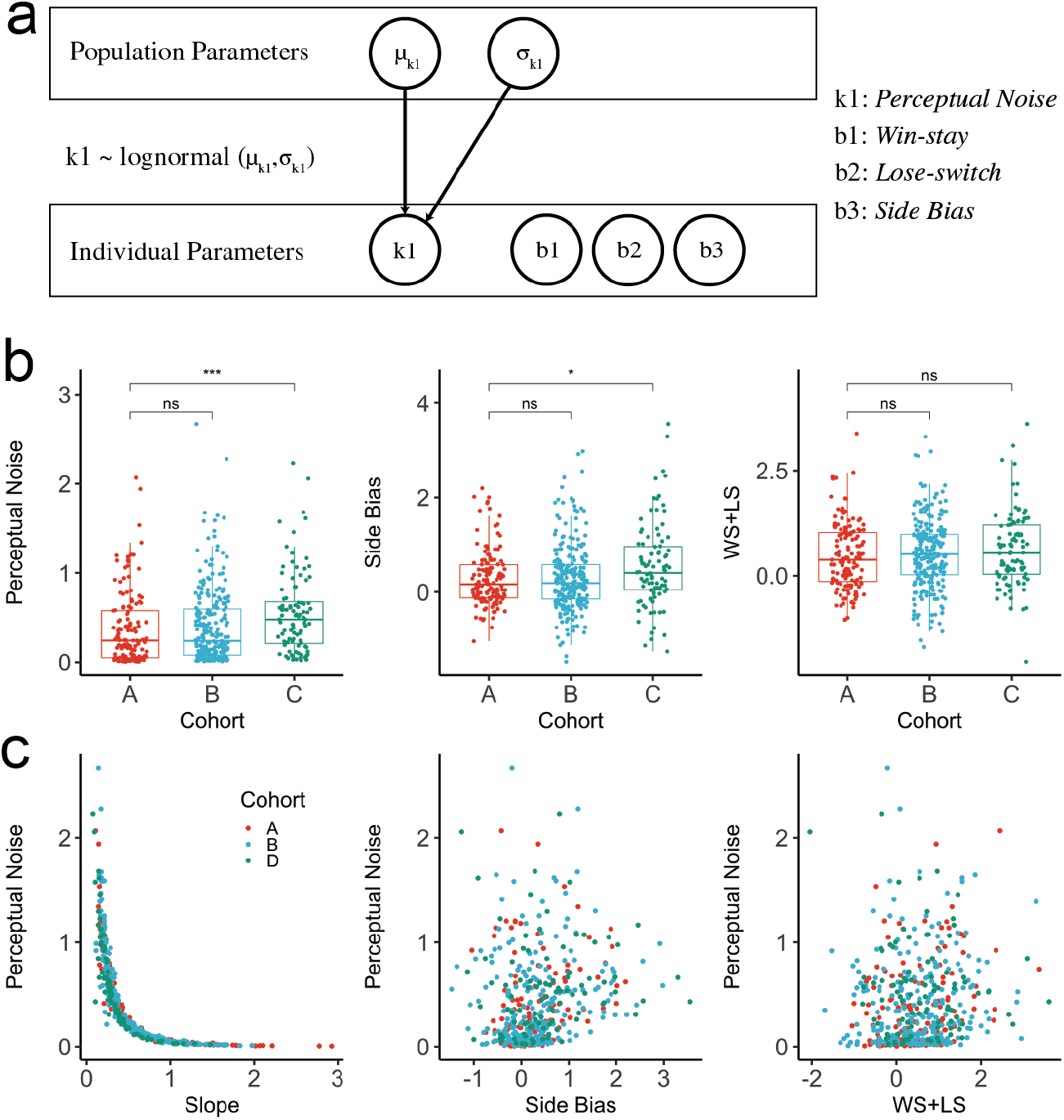
Heterogeneity of behavioral strategies across individuals. (a) A Hierarchical Bayesian Model (HBM) fit to the choice data returned a perceptual noise scaling factor at both the population (µ_k1_, σ_k1_) and individual level (k1), as well as side bias and win-stay lose-switch parameters at the individual level. This schematic shows a subset of the parameters (See **Supplementary Figure 3** for the detailed Model) (b) Pairwise comparisons of the perceptual noise parameter distributions for all participants who passed the selection criterion in Cohorts A vs B (p = 0.27), and A vs C (p = 0.00025) using Mann-Whitney U Test. Pairwise comparisons of side bias parameter distributions for all participants who passed the selection criterion in Cohorts A vs B (p = 0.896), and A vs C (p = 0.012) using Mann-Whitney U Test. Pairwise comparisons of win-stay lose-switch parameter distributions for all participants who passed the selection criterion in Cohorts A vs B (p = 0.26), and A vs C (p = 0.16) using Mann-Whitney U Test. (c) Scatterplots showing the relationship between perceptual noise from the HBM model and psychometric slope from the previously described generalized linear model, perceptual noise and side bias from the HBM, and perceptual noise and win-stay lose-switch from the HBM for each individual.

The HBM revealed extensive heterogeneity in the magnitude of perceptual noise, bias and trial history effects across individuals. In contrast, the differences among cohorts were more modest (**Figure 5b,c**). Pairwise comparison of Cohorts A and B revealed no significant difference in the distribution of perceptual noise (p = 0.27), side bias (p=0.896), or win-stay lose-switch (p=0.26). We did identify a significant difference between Cohorts A and B in an initial noise parameter k0 (p<0.0001, Mann Whitney U Test) fit to the population level (**Supplementary Figure 3**). This parameter served to account for additional sources of errors that were independent of evidence magnitude and trial history, yet to be explored and characterized. We therefore refrained from trying to interpret what specific suboptimality in decision making this parameter was correlated with.

Interestingly, although perceptual noise strongly correlated with the slope of the psychometric function (linear regression with log transformed data, adjusted R-squared = 0.9515), there was no significant correlation between bias and perceptual noise (linear regression, adjusted R-squared = -0.00156) or trial history effects and perceptual noise (linear regression, adjusted R-squared = -0.0014), suggesting that participant performance can vary independently across different behavioral dimensions. Together these results indicate that this approach can be used to evaluate behavioral strategies in evidence accumulation across large numbers of participants using a limited number of trials.

## DISCUSSION

In this study, our goal was to evaluate perceptual decision making using a novel online game inspired by pulse-based accumulation tasks. We used the game to collect data from participants who received full feedback and instructions and analyzed their overall performance, bias, and perceptual uncertainty across all trials, as well as their accuracy across time. When comparing these participants to those who were given the same task structure but trained only on feedback, we found no significant difference between the two cohorts across multiple chosen criteria. Critically, performance across both cohorts was best fit by models in which perceptual uncertainty scaled non-linearly with the strength of the evidence. These results challenge the prevailing diffusion models widely used to describe evidence accumulation in humans, and suggest that a gamified pulse-based accumulation task utilizing operant training may be a useful platform for large-scale acquisition of human perceptual decision making and rule learning data.

### Scalar Variability in Pulse-based Evidence Accumulation

The perceptual judgments of participants performing continuous and pulse-based accumulation tasks are frequently modeled using integration-to-bound models, such as the drift-diffusion model. These models typically assume that the brain extracts evidence from the presented stimulus at a constant rate perturbed by noise (diffusion), and accumulates the evidence linearly over time (Ratcliff & McKoon, 2008).

In this study, we found that human behavioral choices were better fit by a model in which noise scaled nonlinearly with the magnitude of the presented evidence. This is inconsistent with the independent linear accumulation of noise assumed by popular accumulation to bound models. However, this observation is consistent with previous findings from studies in rodents performing similar pulse-based evidence accumulation tasks (Scott et al., 2015, Koay et al., 2020). This nonlinear relationship between perceptual noise and the magnitude of the stimulus is also similar to scalar variability, which has been observed in numerical cognition, timing and other magnitude descrimination studies (Fernandes & Church, 1982; Gallistel & Gelman, 2000; Gibbon, 1977; Mechner, 1958; Nieder & Dehaene, 2009). Given these similarities some researchers have suggested the possibility of shared mechanisms between evidence accumulation and the estimation of number, time and magnitude in general (Gebuis et al., 2017; Howard & Hasselmo, 2020; Pirrone et al., 2022; Simen et al., 2011, 2016).

Extensions of the DDM have been designed to model performance in numerosity and magnitude comparison tasks (Kang & Ratcliff, 2020; Ratcliff et al., 2018; Ratcliff & McKoon, 2018, 2020; Teodorescu et al., 2016). These models assume that noise scales nonlinearly with the magnitude of the stimulus presented at a single time point. However, they also assume that noise across timepoints is independent of the accumulator value (i.e the decision variable), similar to traditional DDMs. Therefore it is unclear whether these models account for superlinear noise accumulation observed when evidence is presented in a stream of uniform pulses as is described in this work.

### Comparison between humans and rodents on pulse based tasks

Our online game provides a valuable behavioral assessment platform to examine perceptual decision making in humans, and enables a direct comparison between humans and rats. Previous studies have compared the performance of humans and rats to an auditory pulse-based task (Brunton et al., 2013). Human and rat performance was evaluated using a sequential sampling model similar to the DDM, and model parameters, such as perceptual noise and the integration time constant of the accumulation could be directly compared.

In this study we extend these results in four ways: First, our game-based approach allowed for a much larger cohort albeit with fewer trials per individual (200 trials in this study vs 1000 in Brunton et al. 2013). The larger participant pool allowed us to observe evolving learning and decision dynamics, and in combination with hierarchical Bayesian analysis recover individual parameters that could be reflective of meaningful variation in perceptual integration across the human population. Second, human participants in our task could be trained solely with experiential feedback, similarly to how rats are trained on this task, allowing for a direct cross-species comparison. Third, we demonstrated that perceptual noise in humans followed the same superlinear relationship with sensory evidence as was previously demonstrated in rats (Scott et al., 2015). Fourth, we observed that humans exhibited several other suboptimalities, with reward location history being an important factor in choices on subsequent trials, adding systematic noise to the decision making process, also similar to what was observed in rats (Scott et al., 2015). Participants tended to repeat choices that resulted in a reward in the previous trial (win-stay), and to switch options after getting a poor outcome (lose-switch). Trial history would impair the decision process since the correct choices were history-independent.

### Suboptimalities in perceptual decision making

The findings in this paper further add to the ongoing debate on whether human perceptual decision are optimal, as is often assumed by the prevalent Bayesian approach to perceptual decision making (Ernst & Banks, 2002; Körding & Wolpert, 2004; Shen & Ma, 2016; Weiss et al., 2002). One of the appeals of the DDM is that it specifies the optimal stopping time for a particular level of accuracy (Bogacz, 2007), and is equivalent to a Bayesian model (Bitzer et al., 2014). However, a growing body of evidence (Keung et al., 2019), including results presented here, indicate that humans exhibit suboptimal perceptual integration (non-linear scaling of perceptual noise) and are heavily influenced by past rewards and side biases, despite knowing the decision rule. Furthermore we show that these biases persist despite knowing reward conditions and task strategy. Together these findings support the incorporation of suboptimalities into computational models of decision making (Rahnev & Denison, 2018).

### Operant training and effects of learning

Operant feedback-based training approaches have a rich history in human studies and provide several advantages to explicit instructions which are traditionally used in human perceptual decision-making studies (Baron & Perone, 1982; Buskist & Miller, 1982; Schwartz & Lacey, 1988). In particular, the feedback-based training pipeline for human participants in the online game opens up several promising future directions to study rule learning. For example, the inclusion of training trials had a dramatic effect on game performance and learning rate. The approach of breaking down complex concepts by providing a sequence of learning steps of increasing difficulty, or creating a curriculum to make learning difficult things easier, is intuitive and prevalent in the modern education system (Metcalfe, 2009).

Work on training artificial neural networks with a curriculum has demonstrated that training examples can yield better generalization (Bengio et al., 2009; Elman, 1993; Krueger & Dayan, 2009). Recently, simulation using artificial neural networks trained with gradient descent has shown that adaptively maintaining an accuracy rate of 85% during training can lead to exponential improvements in the rate of learning, and that this optimal training difficulty might be applicable to biologically plausible neural networks (Wilson et al., 2019). A large and growing body of research has also established the equivalence between gradient descent training and kernel regression in infinite width neural networks (Jacot et al., 2018), and that understanding gradient descent training requires understanding kernel methods (Belkin et al., 2018). The latest theoretical work on learning in kernel regression demonstrated that adding noisy data may impair generalization, leading to non-monotonic learning curves with many peaks (Canatar et al., 2021). Kernel regression comes with an inductive bias that tends to explain observed data with simple functions/stimulus-response mappings, facilitating sample-efficient learning, and biological codes from neural recordings in the mouse’s primary visual cortex seem to exhibit the same property (Bordelon & Pehlevan, 2021). Curriculum training might therefore bias the “neural codes” toward a simple learnable solution and improve the learning rate, but these ideas should be further tested in a laboratory setting with animal models.

In addition to revealing dynamics of learning during the task, our results raise the intriguing possibility that the rule of the game is a strategy that emerges spontaneously, i.e. without shaping or feedback. Evidence for this comes from our observations in participants who played the game without instructions (i.e. Cohorts B, C and D), yet who showed a modest but significant influence of the stimuli on choice even in the first trial, before the participant had received feedback. Since we excluded individuals with previous experience from playing the game, this suggests that the strategy for choosing the side with the greater number of flashes did not require experience in the game. Interestingly as participants received random feedback, they appeared to abandon this strategy and adopt an increasing side bias over the course of a game session.

### Future directions and potential applications

Finally, our online game offers a potential opportunity to evaluate perceptual decision making in minimally verbal human participants. Of particular interest is how evidence accumulation varies across the human population as studies suggest that sensory integration may play a role in conditions such as autism spectrum disorders (ASD; Robertson & Baron-Cohen, 2017). Several studies have successfully demonstrated behavioral modification using reinforcement based strategies in individuals with ASD (Allen et al., 1964; Davidson, 1964; D’Cruz et al., 2013; Jensen & Womack, 1967; Johnson et al., 2006; Lin et al., 2012; Martin et al., 1968; Richard Metz, 1965; Solomon et al., 2015; Wolf et al., 1963). While several models of altered sensory processing have been proposed to account for deficits in perceptual integration in ASD, the underlying cognitive mechanisms are unknown (Robertson & Baron-Cohen, 2017). In addition, minimally verbal ASD participants are traditionally underrepresented in psychophysical studies. The use of our online game with a feedback-based training pipeline will allow us to characterize deficits in perceptual integration across a range of ASD participants and to distinguish among alternative cognitive models.

## METHODS

### Human Participants

Human-participant procedures were approved by the Boston University Internal Review Board; informed consent was obtained from all participants. Participants (n=971) were recruited via Amazon Mechanical Turk. Sample sizes were determined based on the total number of trials collected in previous rodent studies (Scott et al., 2015, 2017). Eligible participants were those living in the United States with 90% or higher approval rates in previous Human Intelligence Tasks (HITs). All participants indicated that they had never played the game before. After consenting to participate, participants were provided with a link to the online game. After completing 200 trials in 20 minutes or less, participants were provided a unique code which they could then enter to the survey on Mechanical Turk to confirm task completion and receive payment.

### Game Design

Each trial of the game consisted of a sequence of states. A trial started in an initial state in which the player was free to move the agent and explore the virtual environment. Once the agent contacted the treasure chest, the task transitioned to the cue state. In this state, a sequence of flashes from two stars located adjacent to each door was presented. On some trials, the number and timing of flashes was fixed (such as during a training trial, or during trials in which some flash combinations were upsampled to fill the stimulus heatmap in **Figure 1c**) (**Supplementary Figure 1**). In other trials, the flash counts were determined pseudorandomly according to a Poisson process using a generative probability of 30% (for one side) and 70% (for the other side) in each of 10 time bins (**Supplementary Figure 1**). Flash duration was 20ms followed by a 230ms delay. Up to 10 flashes total could be shown on each side, amounting to a fixed 2.5s interval in the cue state. After the cue state, the player was transitioned to the choice state, in which the treasure chest became attached to the agent. The player had to deliver the chest to one of two possible doors, making a decision based on the flashes presented during the previous state. After a choice was made, the player was transitioned to the feedback state, in which their score either increased or remained unchanged. The game then looped back to the initial state.

The software for the game was written in JavaScript using the PixiJS game engine. The code is available to download here: https://github.com/qdo1010/coindm.

### Game Feedback and Instructions

Participants in Cohort A were given full instructions (**provided below**) on the task’s rule as well as full feedback on their performance during the task. Cohort B participants received a brief written overview of the task (see below) but had to learn the rule solely through feedback. In the first 10 trials, pairings of 10-versus-0 or 9-versus-1 flashes were presented to both Cohort A and B. We expected that training trials with large differences would facilitate rule learning. Similarly to Cohort B, Cohort C participants received no instruction and had to learn the rule given only feedback. However, the participants in Cohort C did not receive the first 10 “training” trials and were instead presented immediately with the full flash distribution. Participants in Cohort D received no instruction as well as random, noncontingent feedback on their task performance and were presented with 200 trials of the full flash distribution.

### Behavioral Analysis

Data analysis was performed in R (R Core Team, 2020), and all figures were plotted using the ggplot2 package (Wickham, 2016).

#### Flash timing contribution

Flash timing contribution was quantified using a GLM. The model was implemented in R using the function glm and defined as follows:

choice ∼ binomial(n,π), where n is the number of trials.

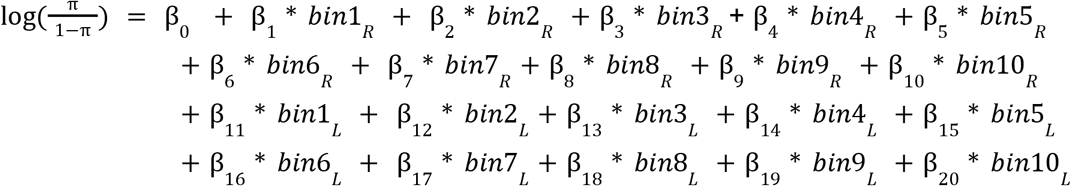

In this model, the probability of going right (π) is determined by the weighted sum of all the flashes. This model has the flexibility to weigh flashes at each time differently. To achieve this, we divide the trial into 10 discrete time bins (250ms each) and the chance of going to the right is regressed against the flashes at each time bin (1 or 0) on both the left and right side. The parameter *bin*1_*L*_ for example is set to 1 when a flash occurs at the first time bin on the left side, and 0 otherwise. Similarly, *bin*1_*R*_ is set to 1 when a flash occurs at the first time bin on the right side, and 0 otherwise. Once fit, the returned weights for each time bin indicate the contribution of individual flashes to the choice of going right.

#### Behavioral generalized linear model

We designed a generalized linear model (GLM) to quantify the effect of the stimuli, previous reward location and side bias on behavior. In this model, the probability of going right (π) depends on a weighted sum of side bias, previous reward (WS), previous omission (LS) and the difference in the number of flashes (*R* − *L*).

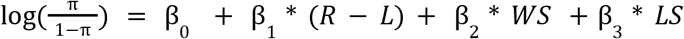

The coefficient β_1_ can be interpreted as the slope of the psychometric function. The WS term was parameterized in the following way: WS = 1 if the previous choice was on the Right Side and was rewarded, WS = -1 if the previous choice was on the Left Side and was rewarded, and WS = 0 otherwise. Similarly, LS = 1 if the previous choice was on the Left Side and was not rewarded, LS = -1 if the previous choice was on the Right Side and was not rewarded, and LS = 0 otherwise.

The generalized linear model was implemented in R using the glm function that assumes a binomial distribution. We fit this model to the pooled choice data for each cohort at single trials and analyzed how the returned parameters changed across time.

#### Analysis of learning rate

The behavioral GLM for each cohort was fit separately for each of 20 bins of 10 contiguous trials. We then fit an exponential learning curve function to the returned psychometric slopes to assess learning across time bins in Cohorts A, B and C.The exponential learning function (Leibowitz et al 2010) was defined as follows:

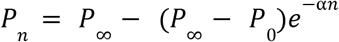

Where n denotes trial number, *P*_*n*_ the fitted performance measured at trial n, *P*_0_ and *P*_∞_ the fitted initial and asymptotic performance, and α the learning rate. the model. *P*_0_, *P*_∞_ and αare free parameters in the model.

The returned psychometric slopes for participants in Cohort D over time were fit with a linear model. Linear models were fit to the bias parameters and win-stay lose-switch parameters for each cohort. We chose linear over exponential models in these cases because the exponential models output errors when fit to the data, and the data seems to follow a linear trend.

#### Signal detection theory model

We implemented a signal detection theory framework to quantify noise in the accumulation process. In this framework, the probability of going to the right is the probability that a random variable Y is greater than 0, where Y is the perceived difference between the total number of right flashes and left flashes, or R and L respectively.

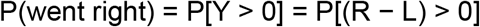

R and L are drawn from a Gaussian distribution N(n_R_, σ_R_) and N(n_L_, σ_L_) respectively. n_L_ and n_R_ are the numbers of flashes presented on each side, whereas σ_L_ and σ_R_ are the associated standard deviations. The difference is a Gaussian where the total mean is the difference between the individual means, while the total variance is the sum of the individual variances. All models were fit using a Maximum Likelihood procedure and simulated annealing was used to solve the optimization problem.

#### Hierarchical Bayesian Analysis

The Hierarchical Bayesian model was written in Stan and fit to the data using the RStan package (Stan Development Team, 2020). The model defines choice at trial i as a random draw from a Bernoulli distribution with probability π_i_, where π_i_ is the probability of going to the right side at that particular trial (**Supplementary Figure 3**). The probability of going to the right side is defined according to the signal detection theory model described previously, with the addition of the win-stay lose-switch parameter and side bias added to the difference in flash number. The perceptual noise parameter k1, was fit on both the population and individual levels. The initial noise parameter k0 from the signal detection theory model was fit only on the population level, and defined as a log normal distribution with mean and standard deviation terms that were shared across the population.

We performed a parameter recovery experiment to verify accurate recovery of k1 (**Supplementary figure 4**). We used the signal detection theory framework to create a cohort of 500 artificial agents whose sensory percept followed the scalar variability model of perceptual noise, with predefined parameters drawn from a log normal distribution. On each trial, we presented these agents with the number of flashes on each side to simulate the post-accumulation step of our human task, and generate choice behavior. We then fit our hierarchical Bayesian model to the resulting synthetic data and compared the returned parameter values with the ground-truth generative parameters. Results confirmed that parameters could be successfully recovered for k1 using as few as 200 trials per participant (r^2^ = 0.77 between ground-truth generative parameter values and recovered values).

### Game instructions and text provided to different cohorts

#### Instructions and text provided to cohort A

“Hello and welcome to Indie and Alone! You will now be competing for a chance to become the next world-renowned archeologist/treasure hunter/action hero. Win points for delivering each treasure chest to the correct door. Pick up the treasure and stars will appear on your Left and Right. Deliver the treasure to the door with more stars. After you make the delivery, a new treasure chest will magically appear. You have 200 chances to win the most points you can. Use the ARROW Keys to move around. Get a High Score. Highest Score is displayed in Red. Don’t Give Up. Be the Very Best that No One Ever Was!”

#### Text Provided to cohorts B, C and D

“Hello and welcome to Indie and Alone! You will now be competing for a chance to become the next world-renowned archeologist/treasure hunter/action hero. Win points for delivering each treasure chest to the correct door. After you make the delivery, a new treasure chest will magically appear. You have 200 chances to win the most points you can. Use the ARROW Keys to move around. Get a High Score. Highest Score is displayed in Red. Don’t Give Up. Be the Very Best that No One Ever Was!”

## Acknowledgments

The authors would like to thank Helen Tager-Flusberg, Rui Cao and members of the Scott lab for helpful discussions. This work was supported by SFARI Human Cognitive and Behavioral Science Award #874568.

## Author Contributions

QD, Concept and Design, Acquisition of Data, Analysis and Interpretation of Data, Drafting or revising the manuscript; GAK, Analysis and Interpretation of Data, Drafting or revising the manuscript; JTM, Analysis and Interpretation of Data, Drafting or revising the manuscript; BBS, Concept and Design, Analysis and Interpretation of Data, Drafting or revising the manuscript.

## SUPPLEMENTS

**Supplementary Figure 1 - Related to Figure 1.**
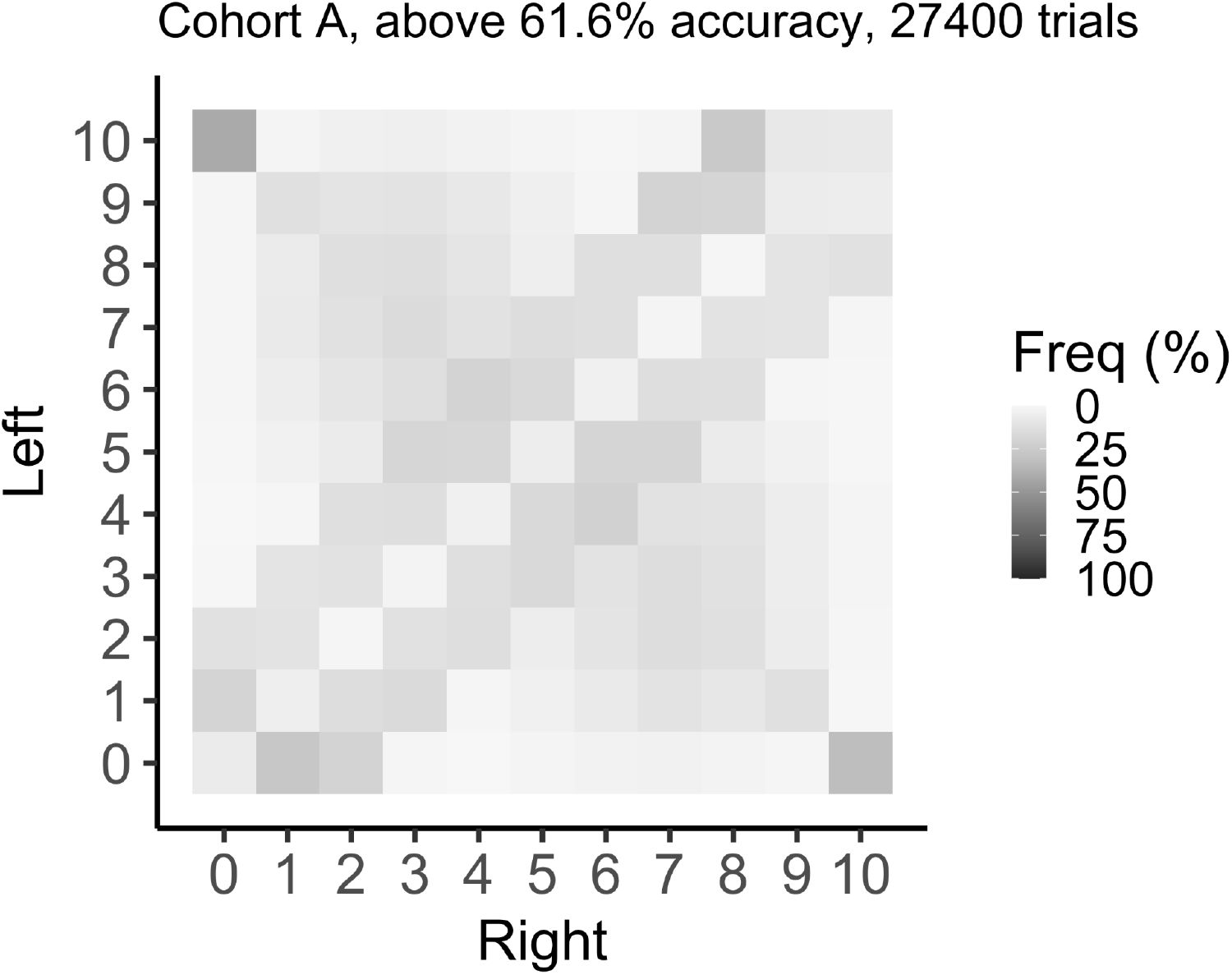
Heatmap showing frequency of flashes pairings reflecting a Poisson sampling process as well as upsampling of difficult pairings (flash difference of 1 and 2).

**Supplementary Figure 2 - Related to Figure 4.**
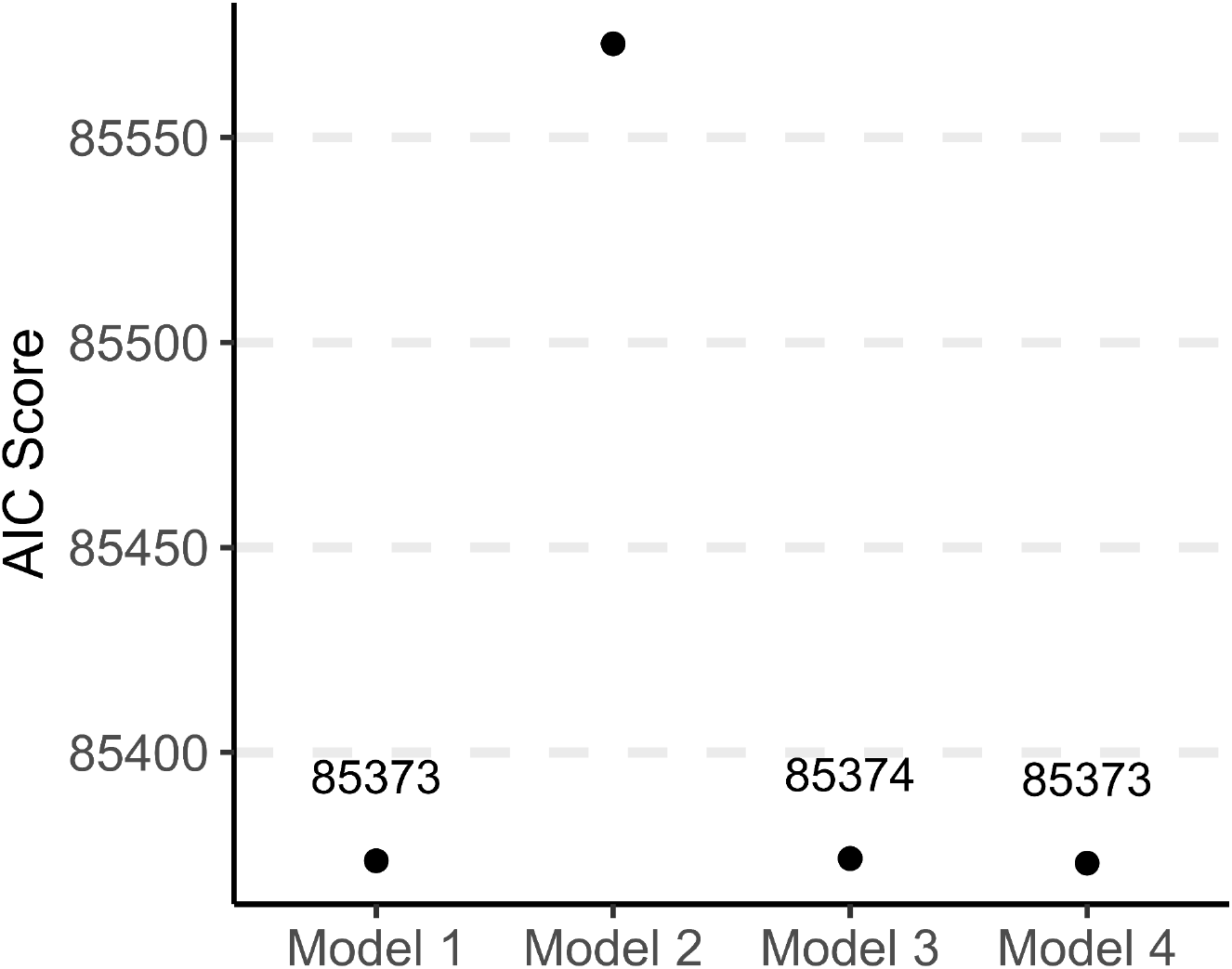
Model comparisons for participants in cohort A, B and C, ignoring the first 30 trials to remove any learning dynamics. Fitted parameters returned by Model 1 (k1 = 1.679, k0 = 0.284), Model 2 (k2=1.58e-05, k0 = 1.85), Model 4 (k1 = 1.328, k2 = 0.255, k0 = 0.209), Model 3 (k = 0.343, k0 = 1.498, ki = 1.903).

**Supplementary Figure 3 - Related to Figure 5.**
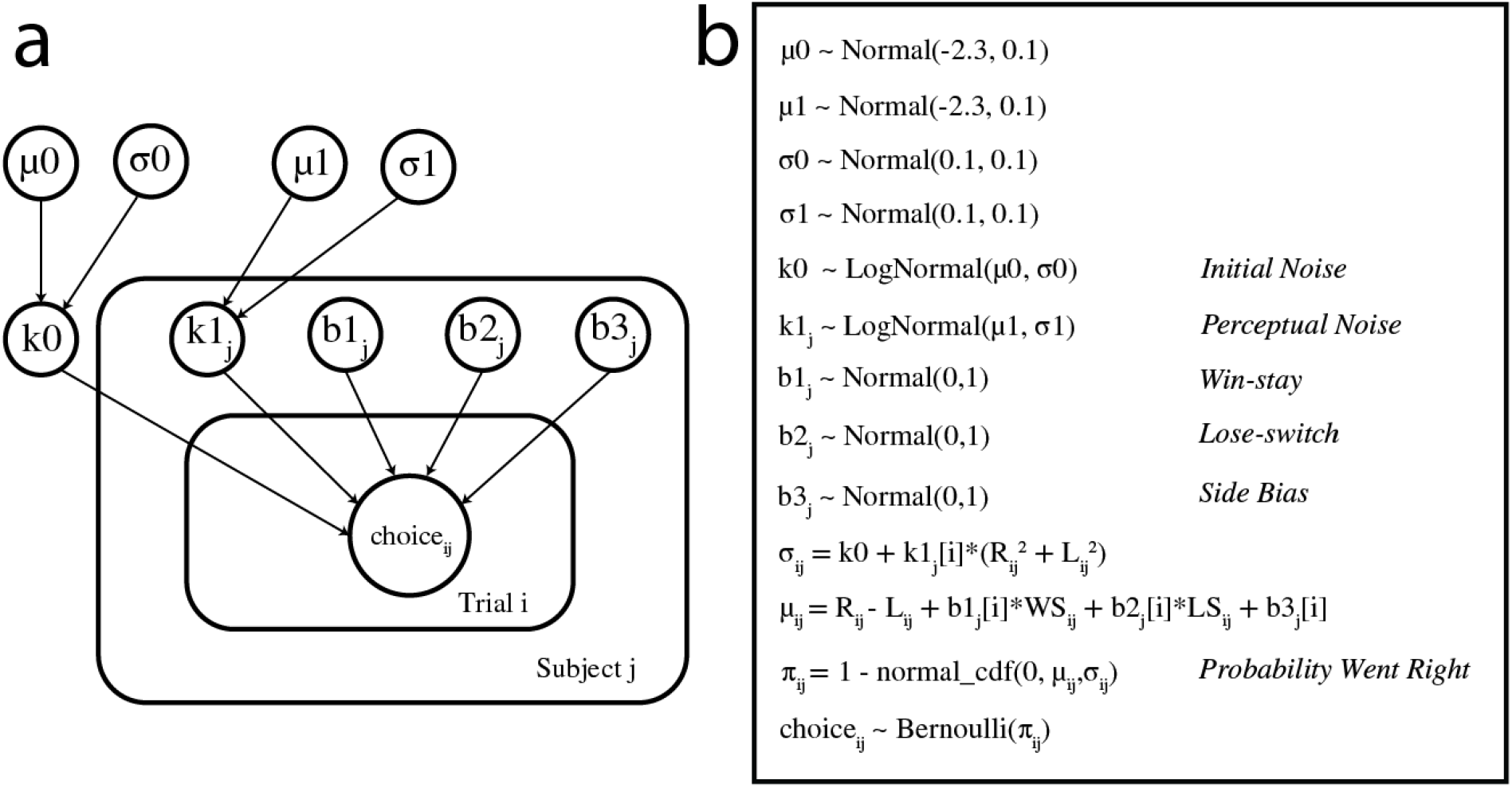
a. Schematic of the Bayesian hierarchical model - signal detection theory model of perceptual uncertainty. b. Fitted parameters and prior distributions for each parameter in the model

**Supplementary Figure 4 - Related to Figure 5.**
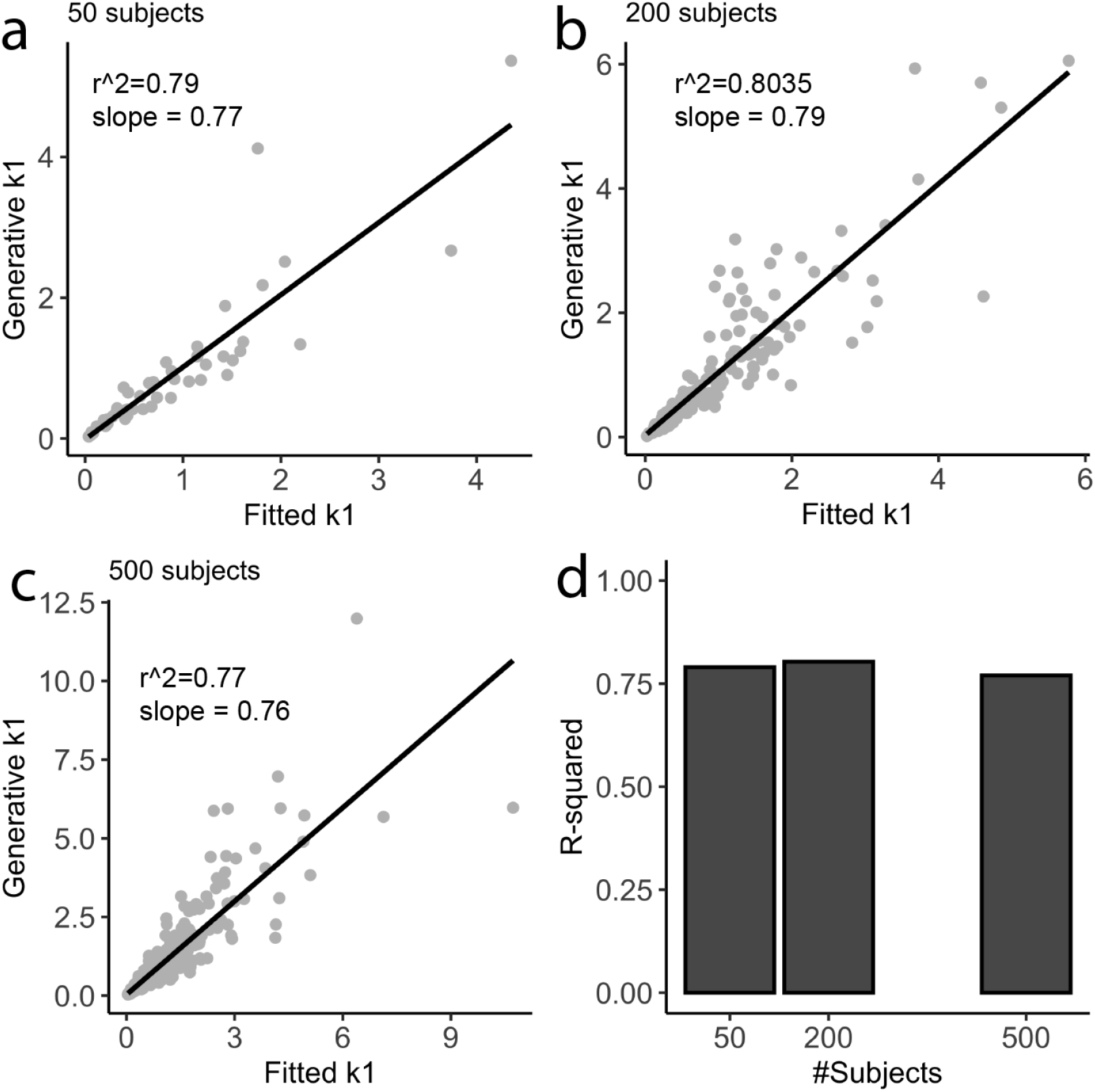
Effect of the number of participants on the accuracy of the recovered parameter using the Bayesian Hierarchical Model. a-c) Fitted k1 from the Hierarchical Model versus Generative k1 for 50 participants, 200 participants and 500 participants doing 200 trials each. d) Comparison of R-squared score with the increasing number of participants.

